# Comparative analysis of inducible promoters in cyanobacteria

**DOI:** 10.1101/757948

**Authors:** Anna Behle, Pia Saake, Ilka M. Axmann

**Affiliations:** Institute for Synthetic Microbiology, Heinrich Heine University Duesseldorf, Duesseldorf, Germany

**Author notes:** Corresponding author; E-Mail | Website: http://www.synmikrobio.hhu.de.

**Keywords:** *Synechocystis*, Inducible promoter, synthetic biology, cyanobacteria, pSHDY

## Abstract

Research in the field of synthetic biology highly depends on efficient, well-characterized promoters. While great progress has been made with other model organisms such as *Escherichia coli*, photosynthetic cyanobacteria still lag behind. Commonly used promoters that have been tested in cyanobacteria show weaker dynamic range or no regulation at all. Alternatives such as native metal-inducible promoters pose the problem of inducer toxicity.

Here, we evaluate four different inducible promoters, both previously published and new, using the modular plasmid pSHDY, in the model cyanobacterium *Synechocystis sp*. PCC 6803 - namely the vanillate-inducible promoter P_vanCC_, the rhamnose-inducible P_rha_, and the aTc-inducible P_L03_, and the Co^2+^-inducible P_*coaT*_. We estimate individual advantages and disadvantages, as well as dynamic range and strength of each promoter in comparison with well-established constitutive systems. We observed a delicate balance between transcription factor toxicity and sufficient expression to obtain a dose-dependent response to the inducer. In summary, we expand the current understanding and employability of inducible promoters in order to facilitate the construction of more complex regulatory synthetic networks, as well as more complicated biotechnological pathways for cyanobacteria.

## Introduction

Cyanobacteria are versatile photoautotrophic organisms that are becoming more and more interesting for various research applications. Due to their ability to fix carbon photosynthetically, they are emerging as promising candidates for the biotechnological production of different compounds, including biofuels^1^ and complex secondary metabolites ^2^.

Their ancestral relation to today’s plant chloroplasts makes them important model organisms for foundational research in the field of photosynthesis^3^. In addition to this, many cyanobacteria are naturally competent and possess the ability to incorporate free DNA into their genomes as well as receive conjugative plasmids, making them attractive from a genetic engineering perspective^4^.

In recent years, more and more tools have been developed and characterized for diverse cyanobacterial species^5^. This includes well-studied model organisms such as *Synechocystis sp*. PCC 6803^6^, *Synechococcus elongatus* PCC 7942^7,8^ and *Anabaena sp*. PCC 7120^9^, but also some more recently discovered, fast-growing strains, such as *Synechococcus elongatus* UTEX 2973^10^ or *Synechococcus sp*. PCC 7002^11^. One challenging aspect when applying promoters previously established in *Escherichia coli* is the difference in RNA polymerase architecture, which results in different binding affinities and overall responses to promoter and operator regions^12^.

Nevertheless, a range of promoters, both constitutive^6,10^ and inducible^13^, has been engineered and successfully implemented in *Synechocystis sp*. PCC 6803 (referred to as *Synechocystis* hereafter). These publications tend to either focus on a single promoter construct with detailed work on sequence variations, or have a different core angle such as metabolic engineering. For the purpose of engineering more extensive synthetic regulatory networks, the availability of multiple differentially regulated promoters or regulatory building blocks is essential; for comparability, they should be characterized in a way that one can easily choose from depending on the application. In order to efficiently fine-tune and optimize more complex systems which combine transcriptional and translational output, fundamental evaluation of precise expression dynamics and strength in the context of a range of constitutive promoters is highly desirable.

One of the most standard inducible promoter systems, which is based on the *lac*-operon from *E. coli* and is inducible by the lactose analog IPTG, has been tested and implemented in some cyanobacterial species. The P_trc_ promoter, for example, performs well in *S. elongatus* and is commonly used in many applications^14^. However, efforts to implement similar constructs in *Synechocystis* have mostly failed, resulting in either extremely leaky expression under non-induced conditions or little to no regulation at all^15^. For example, Camsund *et al.* 2014 investigated sequence-specific repression patterns in *Synechocystis*^16^. They reported a 2.3-fold induction ratio for the original P_trc_ promoter, stating that this was likely due to insufficient levels of the repressor *lacI* in the cell, and furthermore that previous success in *S. elongatus* was likely a result of higher expression of *lacI*. Albers *et al.* 2015 investigated different IPTG-inducible constructs by modification of the gap between the sigma factor binding sites^17^. They placed *lacI* under the control of P_*sigA*_, which promotes expression of the housekeeping sigma factor *sigA* and therefore assures stable, strong expression of the repressor. For their promoter construct P_sca6-2_, they were able to show approximately 10-fold induction ratios.

Another well-characterized promoter in *Synechocystis* is the aTc (anhydrotetracycline)-inducible, *tetR*-regulated system. Huang *et al.* 2013 constructed a library by altering the region downstream of the −10 promoter region^18^. They reported induction ratios of up to 239 for their best performing promoter, P_L03_, under LAHG (light-activated heterotrophic growth) conditions. This promoter suite was also successfully implemented by Yao *et al.* 2016 for dCas9-mediated gene repression, although they reported better success with the weaker, more tightly repressed P_L22_ due to leaky expression of dCas9 from P_L03_^19^. Unfortunately, an issue with aTc in general is the fact that it is light-degradable, making its performance difficult to predict and also preventing it from stable expression under photoautotrophic growth conditions, as are common for most studies focusing on cyanobacteria.

A third system which was reported for *Synechocystis* by Kelly *et al.* in 2018 is the L-rhamnose-inducible promoter P_*rha*_, which is regulated by the transcriptional activator *rhaS*^20^. They thoroughly investigated this promoter under different light and nutrient conditions and found it to be tightly repressed under non-induced conditions, with a linear response upon induction and a good dynamic range, in addition to the inducer, L-rhamnose, being non-toxic to and non-metabolizable by the cells. To date, this is the best working promoter system in *Synechocystis* in terms of both performance and inducer characteristics.

A general issue when selecting promoters for different applications is the data reproducibility. Depending on factors like the choice of measurement methods, (reporter) genes, RBS / 5’UTR or growth conditions, effects on mRNA stability, fold activation or promoter strength may strongly differ between publications^21–23^. While it remains true that each lab should reproduce measurements under their own conditions to ensure reproducibility, an evaluation of constructs in a side-by-side manner using comparable genetic elements and culturing conditions can be helpful in choosing a suitable promoter to begin with.

In contrast to cyanobacteria, there has been ongoing, successful work published for more accessible model organisms such as *E. coli*. For example, Ruegg *et al.* reported the optimization of a promoter system in *E. coli*, previously identified in *Enterobacter lignolyticus*^24^, which responds to a variety of cationic dyes at very low, non-toxic concentrations, including the cheap inducer compound crystal violet, for which they report a dynamic range of four orders of magnitude.

Another recent publication focused on optimization of parameters such as binding of the transcription factor to the operator, full repression under non-induced conditions, and elimination of cross-talk using a two-phase directed evolution approach^25^. Here, a positive selection process involving expression of DNA polymerase was combined with a negative selection involving the toxic expression of a mutant aminoacyl tRNA-synthetase. This yielded 12 highly optimized promoter/sensor pairs, including a vanillate-inducible system originating from *Caulobacter crescentus*.

In this work, we constructed and investigated a comprehensive, comparative library of available different inducible promoters by evaluating them in the same genetic architecture, using the modular plasmid pSHDY. Alongside established aTc-, L-rhamnose-, and Co^2+^-inducible systems, we also present the newly tested vanillate-inducible promoter system.

Finally, we estimated individual promoter performance in a controlled setting for various downstream applications.

## Results and Discussion

### Design framework of all promoter constructs tested in *Synechocystis*

In order to assay each promoter while ensuring comparability/reproducibility, a suitable reporter system was required. We considered a vector with two spatially separated cloning sites, in which the reporter construct comprising promoter, RBS and reporter gene could be located distantly from the repressor/activator and could easily be switched out. For this reason, we constructed pSHDY, a conjugative shuttle vector based on pVZ321^26^, but much more suitable for cloning due to multiple restriction sites flanking the antibiotic resistance cassettes. In addition, pSHDY also contains the *mobA*Y25F point mutation investigated by Taton *et al.*, which leads to an increase in supercoiled plasmid and therefore more efficient downstream cloning applications such as restriction digest^27^.

The basic pSHDY cloning vector contains a total of three antibiotic resistance cassettes, chloramphenicol and kanamycin, which are flanked by two independent cloning sites termed the BioBrick and the NeoBrick site, respectively, and a spectinomycin resistance separating the two (*Fig. 1A*). For the purpose of comparability, we cloned each promoter/reporter construct into the BioBrick site, while keeping the corresponding repressor constructs in the NeoBrick site (*Fig. 1B*).

**Fig. 1:**
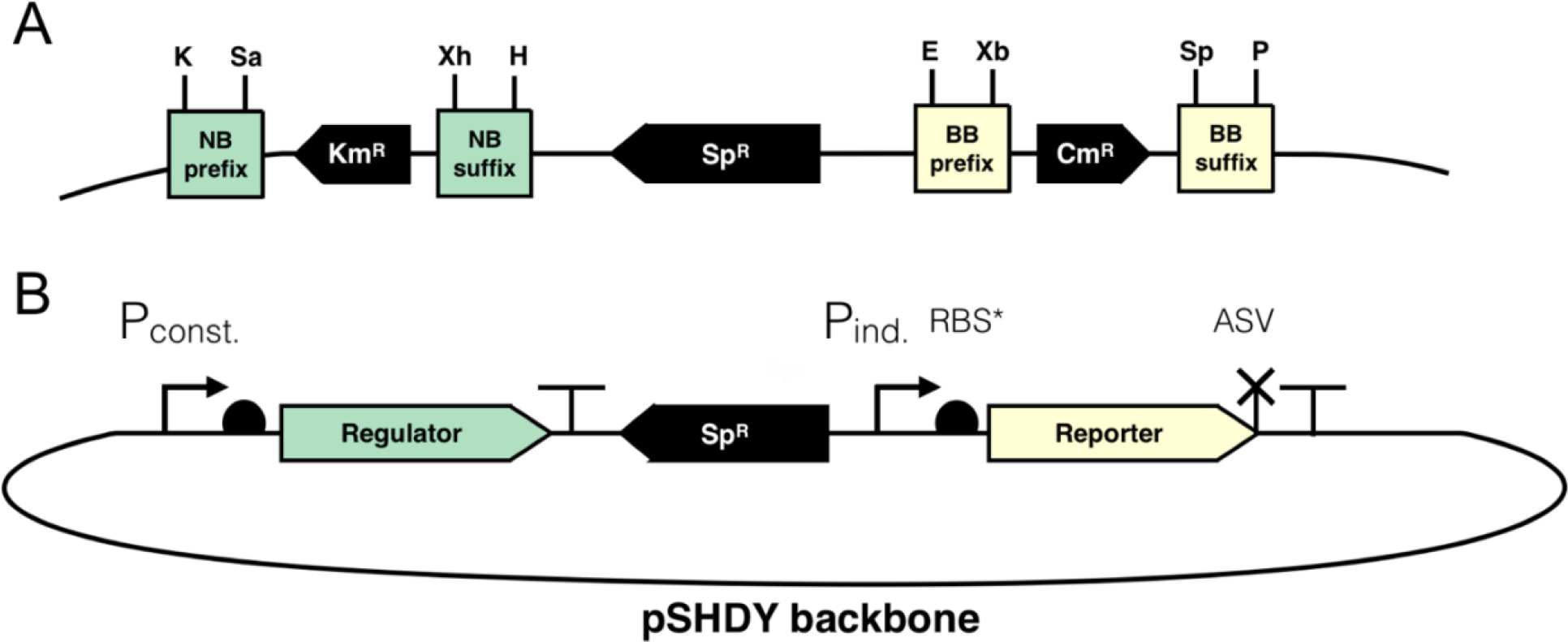
Genetic composition of the different promoter and sensor constructs measured in this work. **A:** Detailed overview of the two modular cloning sites, the NeoBrick (NB) shown in green and the BioBrick (BB) sites shown in yellow. Restriction site abbreviations: K: *Kpn*I; Sa: *Sal*I; Xh: *Xho*I; H: *Hind*III; E: *Eco*RI; Xb: *Xba*I; Sp: *Spe*I; P: *Pst*I **B:** Overview of the plasmid composition of reporter constructs used in this work. P_const_: Constitutive promoter. Sp^R^: Spectinomycin resistance. P_ind_: inducible promoter. ASV: *ssrA*-based ASV-degradation tag.

The promoter/reporter devices were constructed in a comparable manner. For the reporter, we chose mVenus, an eYFP variant with enhanced brightness^28^. The gene was further codon-optimized for *Synechocystis* and a C-terminal *ssrA*-based ASV degradation tag for moderate protein turnover^29^ was added to prevent measuring stable protein instead of inducer-dependent expression. For the RBS, we used the established synthetic RBS*^30^, which was shown to perform well in *Synechocystis* on multiple occasions, except in the native promoter constructs, P_*coaT*_, as well as P_cpc560_, which has been reported to require its native RBS for maximum strength^10^. In the case of P_vanCC_, the riboJ insulator was kept in the 5’UTR as was constructed in the original publication^25^.

Table **1** contains a basic description of all promoters tested in this work, while detailed descriptions and sequences of each promoter construct can be found in the Supplementary information (Table S1).

**Table 1:**
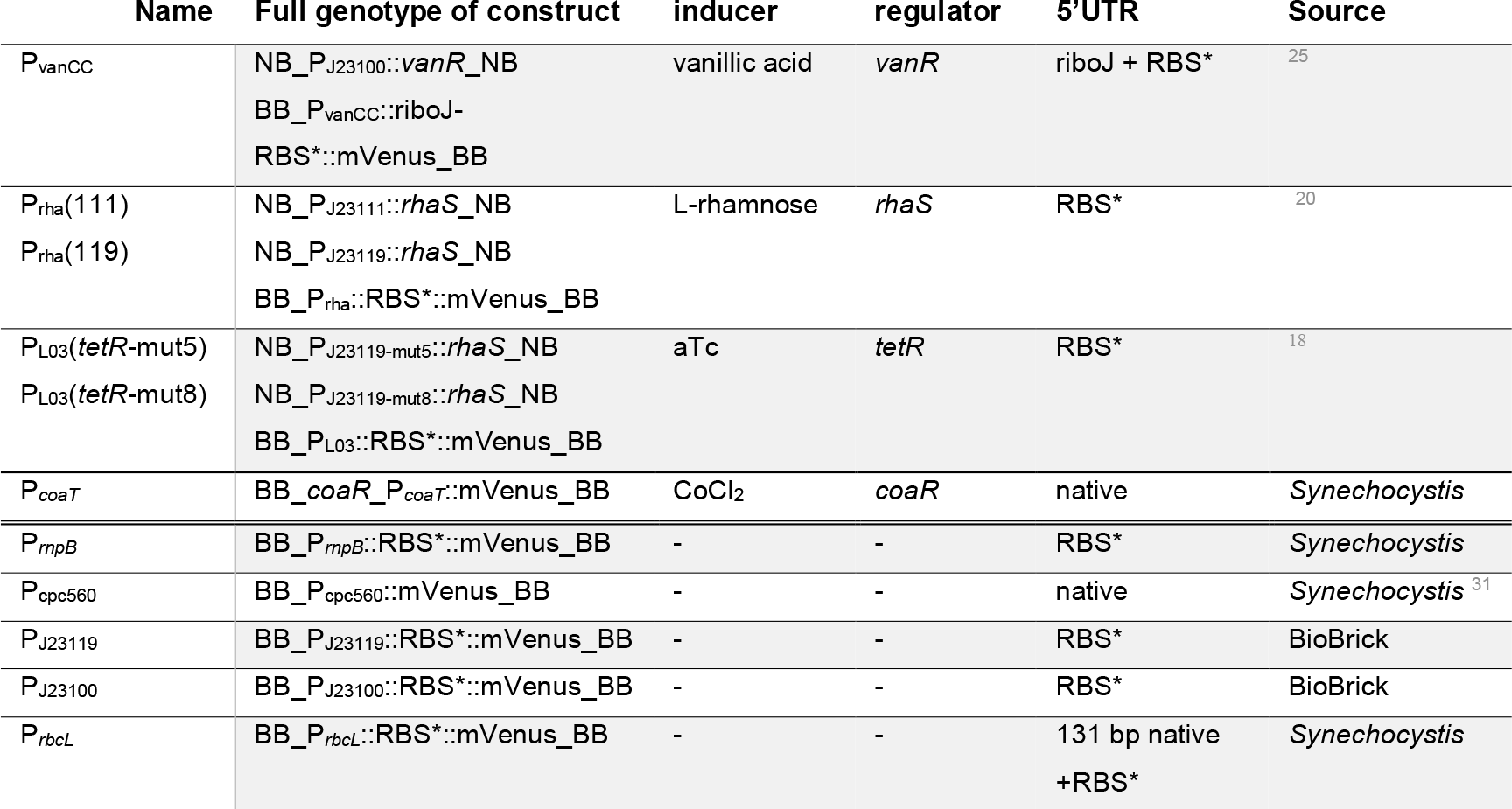
Overview of promoter constructs tested in this work. Inducible promoters are shown above, constitutive promoters below the double line.

Another consideration was encoding the repressor constructs on the genome and the promoter constructs on a plasmid, but we decided against it for two reasons.

Firstly, the copy number of the *Synechocystis* genome can fluctuate depending on different conditions such as growth phase, light intensity or nutrient availability, potentially resulting in different repressor copy numbers and subsequent strength of gene repression. In contrast, plasmid copy number is more stringently regulated within the cell, leading to more consistent results^32^. This also relates to the fact that expression may vary depending on the genomic context. Since different working groups have been using different genomic integration sites, data may not be directly reproducible.

Secondly, it takes longer to generate fully segregated genomic mutants, extending the amount of time between conceiving a project and measuring the data, further complicating rapid genetic screens.

Therefore, we determined a plasmid-encoded reporter system to be the best option.

### Introducing the vanillate-inducible promoter P_vanCC_ in *Synechocystis*

While there have been publications on vanillate inducible systems, mainly in α-proteobacteria^33^, but also in *E. coli*^25^, to the best of our knowledge, this is the first detailed, dose-dependent promoter study in *Synechocystis*. One publication focusing on NOT-gates in *S. elongatus* PCC 7942 used the promoter/repressor pair *vanR*/P_*vanA*_ from *Corynebacter glutamicum*^34^. There, the regulation of the promoter-sensor pair was successfully implemented independently of the inducer vanillate. However, vanillate-dependent induction was not further investigated.

We ultimately chose to evaluate *vanR*/P_vanCC_ from the recent publication by Meyer *et al.*, 2018^25^, in *Synechocystis*. Here, the promoter/repressor pair *vanR*/P_vanCC_ from *Caulobacter crescentus* was rationally designed and then further optimized via directed evolution for *E. coli*, yielding a vanillate sensor with both improved dynamic range, as well as lower cross-reactivity.

We chose the weak constitutive promoter P_J23100_ from the Anderson library and the published van3 RBS to control *vanR*, shown schematically in Fig. 2A. The van3-*vanR* fusion was amplified from sAJM.1504, the Marionette-Clo strain (addgene ID 108251). For P_vanCC_, we amplified the original promoter construct, including the riboJ insulator, from pAJM.714 (addgene ID 108515), but replaced the RBS with the synthetic RBS*. Detailed descriptions and sequences are provided in the supplementary information (Table S1). The conjugative plasmid containing P_J23100_∷*vanR* and P_vanCC_∷mVenus were transferred into *Synechocystis* via conjugation. Transconjugants were validated, cultured, induced and mVenus fluorescence, as well as the optical density at 750 nm, was measured in a microplate reader. An empty vector control (EVC) was included for each concentration. 24h post-induction, we observed a linear dose-response to vanillate, which saturated at 1 mM (Fig. 2B). Furthermore, under non-induced conditions, the promoter remained tightly repressed, reaching the same autofluorescence levels observed in the empty vector control.

**Fig. 2.**
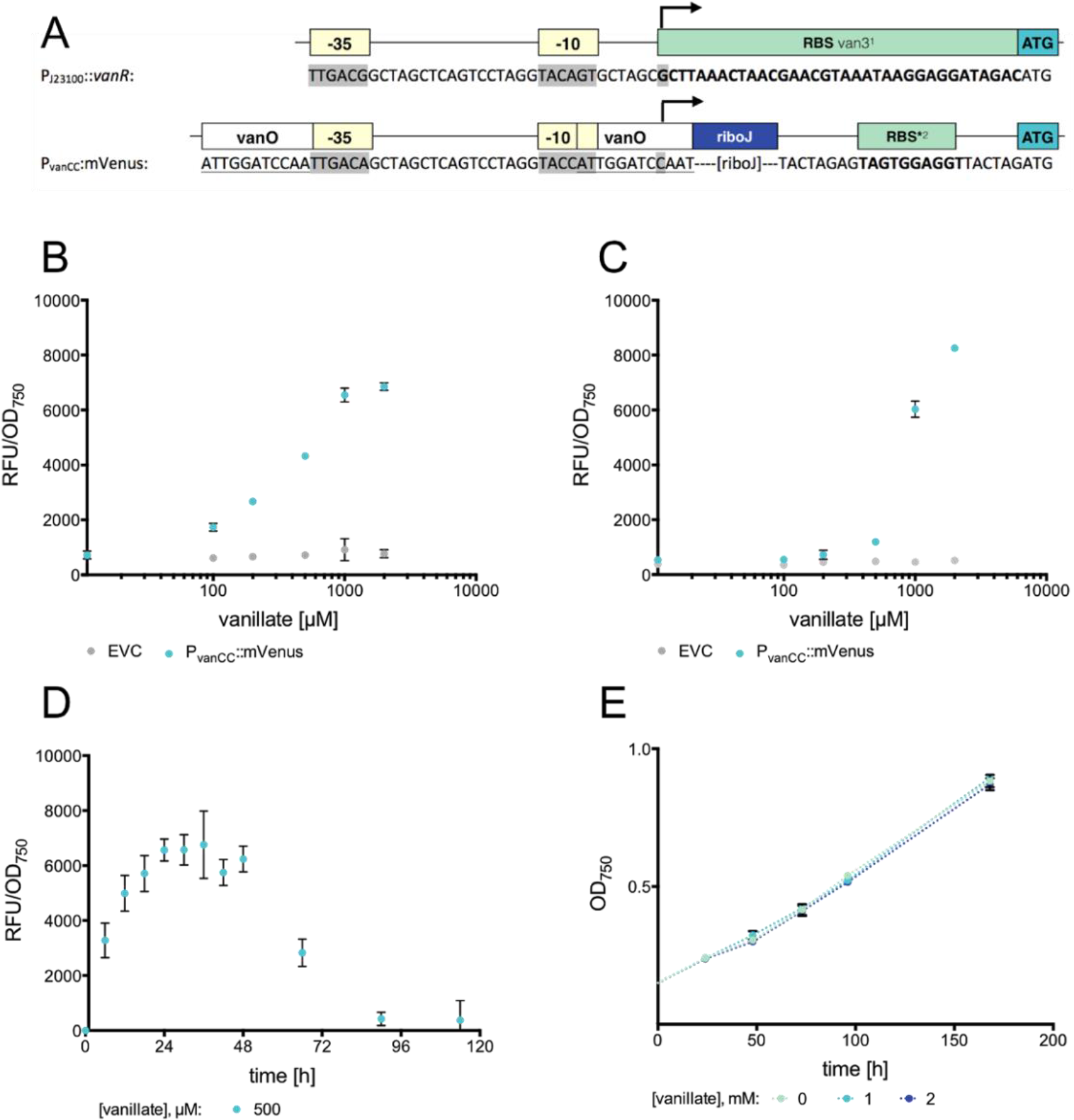
Dose-dependent response of the vanillate-inducible promoter in *Synechocystis*. **A:** Schematic overview of genetic construct used. Top: Genetic composition of regulator. Bottom: Genetic composition of regulated promoter. −10, −35 and +1 are highlighted in grey; RBS is shown in bold. Operator regions are underlined. **B:** Dose-response of the vanillate-inducible promoter P_vanCC_ to different concentrations of vanillate after 24h **C:** Dose-response of the vanillate-inducible promoter P_vanCC_ to different concentrations of vanillate after 48h **D:** Response of P_vanCC_∷mVenus to 500 μM vanillate over time. **E:** Growth of WT *Synechocystis* in different vanillate concentrations. Three biological replicates were cultured in BG11 + vanillate and fluorescence and optical density was measured in a microplate reader. ^1^van3 RBS ^25^; ^2^ RBS* from ^30^

After 48h, a decrease in fluorescence to a fraction of that after 24h could be observed at lower concentrations (100-500 μM), while fluorescence increased or remained at a similar level at saturating concentrations of 1-2 mM (Fig. 2C). While there is no evidence of light-mediated degradation of vanillate, it is an intermediate in the biochemical degradation of lignin^35^, so we hypothesized that vanillate might be degraded by *Synechocystis* after longer periods of time. We therefore chose to investigate mVenus fluorescence over time. To account for possible inducer degradation, we chose a vanillate concentration of 500 μM, which was below saturation of expression and at which concentration a decrease in fluorescence was observable.

After induction, cultures were measured every 6h. To account for effects caused by cell density, an aliquot of each culture was sampled and cell density was adjusted to the start OD_750_ of 0,25 prior to each measurement.

P_vanCC_ rapidly responded to vanillate induction, reaching a fluorescence maximum approximately 24h post-induction. This level was maintained until 48h post-induction, after which fluorescence gradually decreased in a linear fashion, reaching autofluorescence levels 90h post-induction (Fig. 2D).

Since vanillate appears to be completely degraded by *Synechocystis* after 90h, we chose to investigate whether it had any effect on its growth. WT *Synechocystis* cells were treated with different concentrations of vanillate, and OD_750_ was monitored over 7 days (Fig. 2E). Interestingly, vanillate had no positive or negative influence on growth of *Synechocystis*. We therefore hypothesize that degradation of vanillate occurs in an unspecific manner, not significantly contributing as a nutrient.

Overall, P_vanCC_ performs well in *Synechocystis* in a dose-dependent manner, showing no toxicity, tight repression and good dynamic range, with a maximum fold-induction of 16× (2 mM vanillate, 48h post-induction).

### The strong rhamnose-inducible promoter can be fine-tuned via activator expression

While the degradation of vanillate shown for P_vanCC_ can be positive for certain applications, it can also be a drawback when a more long-term expression is desired.

Since the P_rha_ promoter published by Kelly *et al.*^20^ showed such promising results both in dynamic range and stability over time, we aimed to reproduce the data. In accordance with our design framework, which allows for modular exchange of parts, we chose to investigate whether L-rhamnose response could be further tuned by fusing two different minimal constitutive promoters upstream of the activator gene *rhaS* - P_J23119_, containing the *E. coli* consensus core elements and reportedly the strongest of the Anderson promoter library, and P_J23111_, which was shown to be approximately half as strong as P_J23119_ in *Synechocystis*^10^ (Fig. 3A). Plasmids containing these fusions, as well as P_rha_∷mVenus, were transferred into *Synechocystis* via conjugation. Transconjugants were validated, cultured, induced and mVenus fluorescence, as well as the OD_750_, was measured in the plate reader. An empty vector control (EVC) was included for each concentration. The strains were termed P_rha_∷mVenus(119) or P_rha_∷mVenus(111), with the number in parentheses corresponding to the respective Anderson promoter number.

**Fig. 3.**
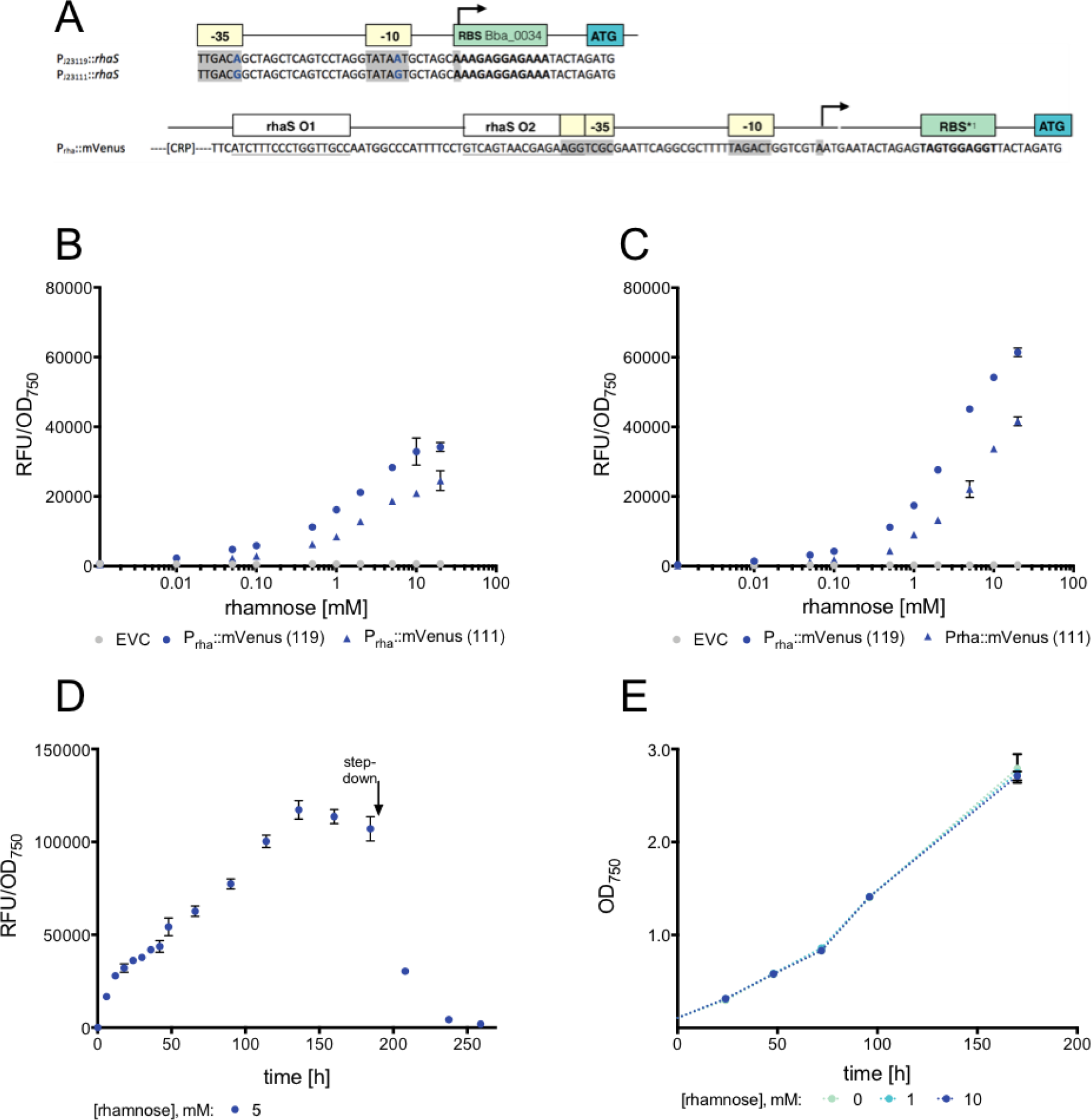
Dose-dependent response of the rhamnose-inducible promoter P_rha_ in *Synechocystis*. **A:** Schematic overview of genetic constructs used. Top: Genetic composition of regulator. Bottom: Genetic composition of regulated promoter −10, −35 and +1 are highlighted in grey; RBS is shown in bold. Operator regions are underlined. **B:** Dose-response of the rhamnose-inducible promoter P_rha_ to different concentrations of aTc after 24h **C:** Dose-response of the rhamnose-inducible promoter P_rha_ to different concentrations of aTc after 48h **D:** Response of P_rha_::mVenus(119) to 5 mM rhamnose over time. OD750 of each sample was adjusted to 0,25 prior to fluorescence measurement. **E:** Growth of WT *Synechocystis* in different rhamnose concentrations. Three biological replicates were cultured in BG11 + inducer and fluorescence and optical density was measured in a microplate reader. ^1^ RBS* from *^30^*

A typical dose-dependent response can be observed in both reporter constructs 24 h post induction, saturating at approximately 10 mM L-rhamnose (Fig. 3B). A maximum fold induction of 55× and 39× is achieved for P_rha_∷mVenus(119) and P_rha_∷mVenus(111) at this concentration, respectively. While Kelly *et al.* don’t specify fold change, a 15× increase under similar conditions can be roughly estimated from their data, indicating an overall improvement of promoter function. We hypothesize that this improvement can be attributed to both an increase of activator levels inside the cell and the use of the well-established RBS* instead of the native *E. coli* RBS, which may increase the maximal exression achievable with P_rha_.

When growing induced cultures for a longer amount of time, the general dose-dependent pattern remained the same for both strains. However, both overall fluorescence intensity, as well as fold induction at 10 mM appeared to further increase over time, up to 165× and 143× after 76 h for P_rha_∷mVenus(119) and P_rha_∷mVenus(111), respectively (Fig. 3C). Therefore, we decided to also evaluate the short- and long-term temporal expression dynamics. To account for possible inducer degradation and reliably assert expression dynamics, we chose a rhamnose concentration of 5 mM, which was below saturation of expression.

After induction, cultures were measured every 6h. To account for effects caused by cell density, an aliquot of each culture was sampled and cell density was adjusted to the start OD_750_ of 0,25 prior to each measurement.

Fluorescence rapidly increased directly after induction. 18h post-induction, this increase became linear. Fluorescence continued to increase linearly until 136h post-induction, after which fluorescence levels remained stable for three more days (Fig. 3D). To investigate whether this was reversible, we performed a step-down by washing the cells twice with BG11 to remove all L-rhamnose from the media. OD_750_ was adjusted to 1.0. Fluorescence rapidly decreased after step-down, reaching pre-induction autofluorescence levels after 3 days.

Finally, we chose to investigate whether L-rhamnose had any effects on *Synechocystis* growth, since some of the concentrations used were higher than previously tested by Kelly *et al.* WT *Synechocystis* cells were treated with different concentrations of L-rhamnose, and OD_750_ was monitored over 7 days (Fig. 3E). Consistent with previous results, L-rhamnose had neither a positive nor a negative effect on *Synechocystis* growth. Moreover, the fluorescence time-course results further support the hypothesis that *Synechocystis* is unable to use L-rhamnose as a carbon source.

As already stated by Kelly *et al.*, P_rha_ performs exceptionally well as an inducible promoter, with a high dynamic range, tight repression, stable expression over at least 7 days, and no toxic effects or metabolization of the inducer.

### The aTc-inducible promoter P_L03_ shows improved function by increasing the protein levels of the *tetR* repressor

Next, we chose to evaluate the P_L03_ promoter published by Huang and Lindblad^18^. Despite their promising results of 300× fold induction this promoter was also reported to lead to leaky expression of dCas9 by Yao *et al*.^19^. For these reasons, we chose multiple different design strategies to possibly reduce leakiness. Firstly, we removed the *ssrA*-based LVA degradation tag to overcome rapid degradation of the repressor protein. Secondly, we chose the strong promoter P_J23119_ in place of P_J23101_ to further increase intracellular TetR. Finally, we applied the same plasmid-based design strategy used for the other promoters to be able to compare the results later on.

Interestingly, we were unable to obtain clones with the expected regulatory sequences upstream of *tetR* planned *in silico*. Instead, each sequenced clone showed point mutations either in the promoter or RBS sequence, suggesting toxicity resulting from excessive expression of *tetR*. Since we preselected clones that showed no fluorescence in *E. coli* for sequencing, indicating tight repression of P_L03_ in *E. coli*, we decided to investigate two of them despite the point mutations. We termed them *tetR*-mut5 and *tetR*-mut8. Fig. 4A highlights the genetic composition of the two mutants compared to the desired construct. Cultures containing the plasmid constructs were treated identically to the ones containing the P_vanCC_ and P_rha_ promoter constructs. For the purpose of employing this promoter in broad, standard applications, we limited our experimental setup to photoautotrophic growth conditions (see the Method section for details), despite Huang *et al.* reporting better results for cultures grown in red light and LAHG.

**Fig. 4.**
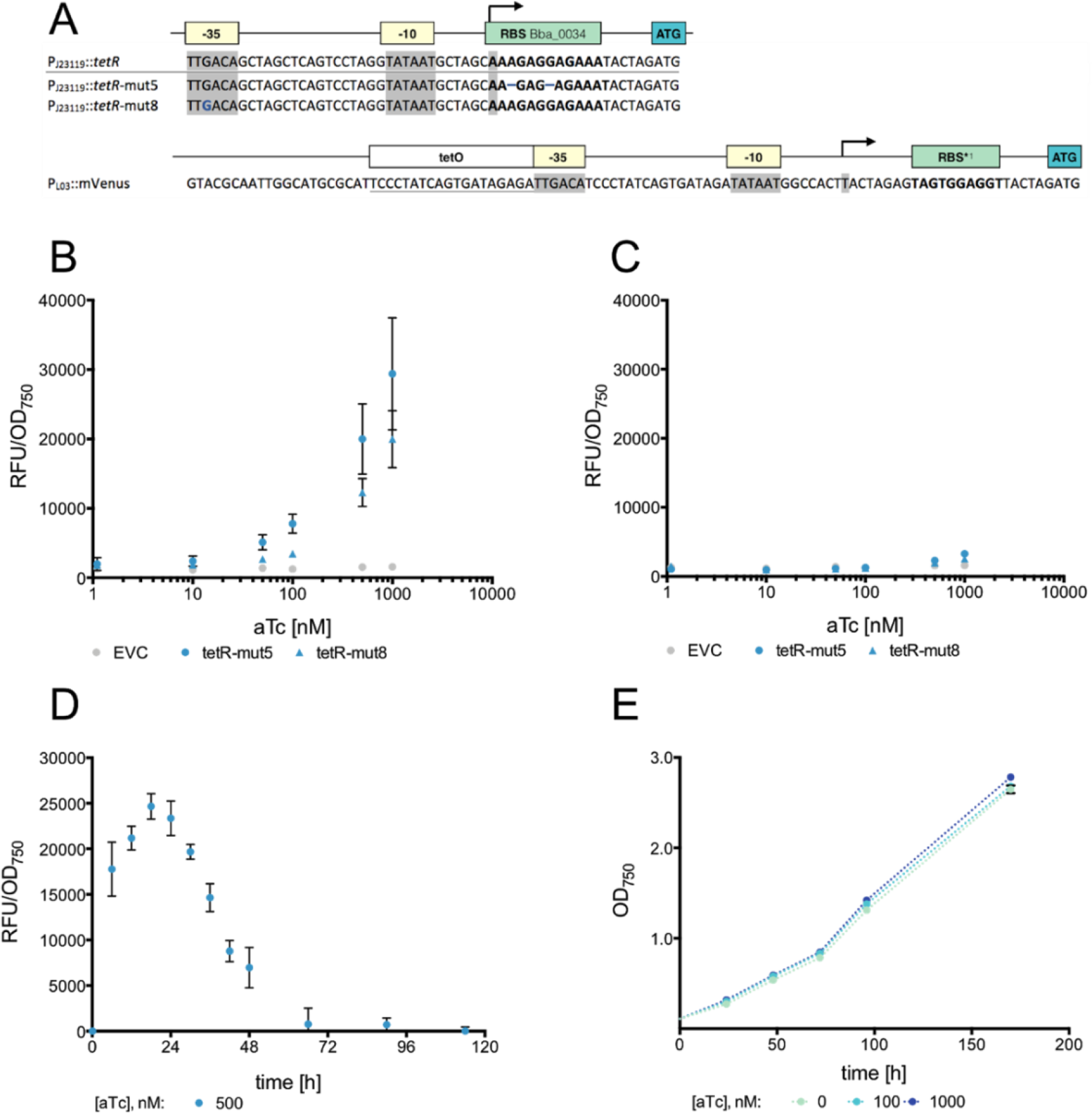
Dose-dependent response of the aTc-inducible promoter P_L03_ in *Synechocystis*. **A:** Schematic overview of the mutant variants with the intended construct (P_J23119_∷*tetR*) as a reference. Top: Genetic composition of regulator. Bottom: Genetic composition of the regulated promoter P_L03_. −10, −35 and +1 are highlighted in grey; RBS is shown in bold. Point mutations / deletions are shown in blue. **B:** Dose-response of the aTc inducible promoter P_L03_ to different concentrations of aTc after 24h **C:** Dose-response of the aTc inducible promoter P_L03_ to different concentrations of aTc after 48h **D:** Growth of *Synechocystis* WT supplemented with different concentrations of aTc. Three biological replicates each were cultured in BG11 and measured in the spectrophotometer. **E:** Response of P_L03_ (*tetR*-mut5) to 500 nM aTc over time. OD_750_ of each sample was adjusted to 0,25 prior to fluorescence measurement. Two and three biological replicates for *tetR*-mut5 and *tetR*-mut8, respectively, were cultured in BG11 + inducer and fluorescence and optical density was measured in a microplate reader. ^1^ RBS* from *^30^*

Fig. 4B shows the dose response of the two mutant constructs 24h post-induction. Interestingly, the dose response assay shows the expected linear aTc-dependent increase of relative fluorescence. The fold change at 1000 nM aTc was lower than for rhamnose with 16-fold and 11-fold for *tetR*-mut5 and *tetR*-mut8, respectively. While the *tetR*-mut8 strain outperforms *tetR*-mut5 both in dynamic range and maximum strength of the promoter in terms of relative fluorescence achieved, it also shows minimally higher leaky expression under uninduced conditions (Fig. 4 B).

The dynamic range of mVenus expression decreased over time; by 48h post-induction, fluorescence had significantly decreased to a fraction of what was measured before (Fig. 4C). We therefore decided to also evaluate the short- and long-term temporal expression dynamics.

To account for possible inducer degradation, we chose an aTc concentration of 500 nM, which was below saturation of expression.

After induction, cultures were measured every 6h. To account for effects caused by cell density, an aliquot of each culture was sampled and cell density was adjusted to the start OD_750_ of 0,25 prior to each measurement.

Consistent with the results observed for P_vanCC_ and P_rha_, fluorescence rapidly increased, reaching a maximum after 18h (Fig. 4D). However, in contrast to P_vanCC_, fluorescence decreased again just as rapidly, reaching autofluorescence levels after 66h. In accordance with published literature, the rapid decrease in fluorescence is most likely a result of light-mediated degradation of aTc. As reported in Huang *et al.*, this promoter likely performs much better under LAHG in darkness or red light. However, for reasons stated earlier, we chose not to further investigate P_L03_ under these conditions.

Since aTc is a derivative of the antibiotic tetracycline, there have been reports on its toxicity in *E. coli* at high concentrations^36^. Thus, we were interested in its effects on the growth of *Synechocystis* WT at the relevant concentrations used for induction of P_L03_.

Interestingly, aTc-treated cells show minimally improved growth compared to untreated cells (Fig. 4E). We attribute this effect to hormesis, a positive effect on growth often observed in bacteria as a result of a global stress response to sublethal concentrations of antibiotics^37^. At concentrations relevant for the induction of P_L03_, aTc appears to have no growth-inhibiting effect on *Synechocystis*.

As previously shown by Huang *et al.*, P_L03_ performs well as an inducible promoter. Providing a suitable intracellular amount of TetR, it shows minimal leakiness and good dynamic range. Especially during the first 24h, it shows rapid, strong induction, making it a suitable tool for applications within this time-frame. Due to the light-sensitive properties of aTc, this promoter may be better suited under red light or darkness for longer term induction experiments. It also may be beneficial for the half-life of aTc to adjust the culture conditions to a higher cell density, thereby preserving the aTc due to shading.

### Evaluating the native Co^2+^-responsive promoter, P_*coaT*_, as an inducible promoter

Next, we decided to investigate a commonly used metal-inducible promoter. Since the most efficient and commonly used Ni^2+^-responsive promoter, P_*nrsB*_, has already been investigated in detail elsewhere^13^, we chose P_*coaT*_. This promoter was successfully used towards small-scale biotechnological production of plant terpenes^38^ and ethylene^39^ in *Synechocystis*, as well as for mimicking a null mutant in the filamentous cyanobacterium *Anabaena sp*. by selectively removing Co^2+^ and Zn^2+^ from the media^40^.

Since the TSS of P_*coaT*_ has, to the best of our knowledge, not been mapped previously, we performed 5’RACE (rapid amplification of cDNA ends) to determine the TSS of P_*coaT*_. Our results indicate at least 3 putative TSS for P_*coaT*_ (Fig. 5A, Fig. S1). We therefore decided to maintain the native promoter+5’UTR architecture, and fused the entire 1195 bp upstream of *coaT*, including the *coaR* repressor, upstream of mVenus.

**Fig. 5.**
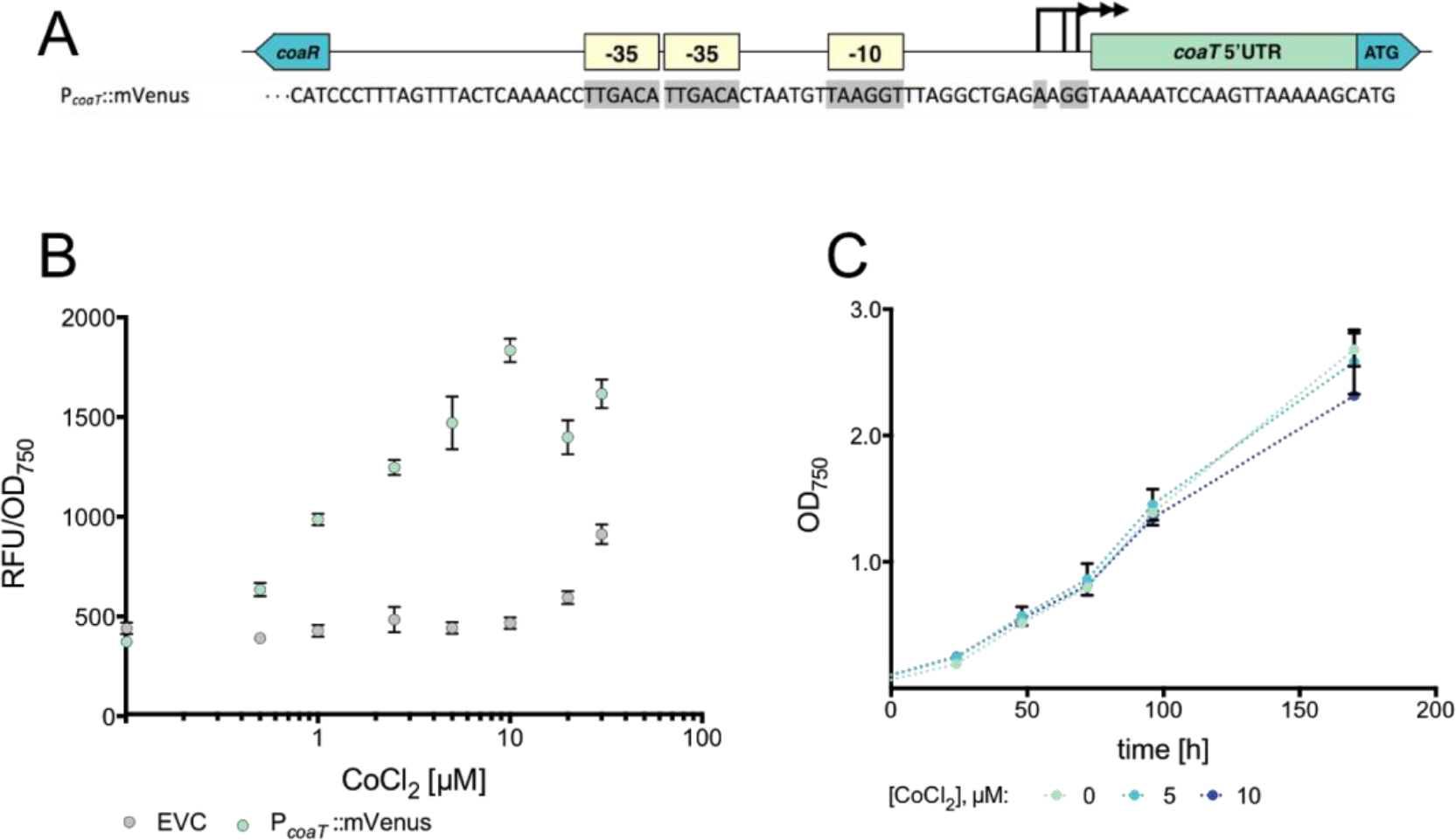
Dose-dependent response of *Synechocystis* to Cobalt. **A:** Schematic overview of genetic construct used. −10, −35 and putative +1 are highlighted in grey. **B:** Dose-response of P_*coaT*_∷mVenus to different concentrations of CoCl_2_ after 48h **C:** Cobalt-dependent growth behavior of WT *Synechocystis* over time. Optical density Three biological replicates were cultured in BG11 + inducer and fluorescence and optical density was measured in a microplate reader.

Upon induction with different concentrations of CoCl_2_, a linear response could be observed up to a concentration of 10 μM (Fig 5B). For higher concentrations, the values measured became erratic for both P_*coaT*_∷mVenus and EVC. This is likely due to toxic effects of Co^2+^ ions.

Upon investigating effects of relevant CoCl_2_ concentrations on growth of WT *Synechocystis*, a slight defect in growth was observed at 10 μM (Fig. 5C). This effect was even stronger in P_*coaT*_∷mVenus (data not shown), which is consistent with previous observations, where it was reported that deletion of *coaT* led to higher cobalt sensitivity^41^. Increasing the amount of CoaR repressor in the cell, as done in this work by expression of an additional copy from a plasmid, likely has the same effect. Moreover, the maximum working concentration of Co^2+^ reported throughout the literature for the P_*coaT*_ promoter is 6 μM^42^ indicating toxic effects at higher concentrations.

More importantly, for complete repression of P_*coaT*_, it is necessary to culture strains in Co^2+^-depleted BG11. Since Co^2+^ ions are required for the synthesis of coenzyme B_12_ in diverse cyanobacteria^43^, this means that complete repression of the promoter may require a defect in growth as a result of nutrient limitation.

When looking into temporal expression dynamics, Englund *et al.* could show a decrease of fluorescence for P_*nrsB*_, due to Ni^2+^ actively being pumped out of the cells ^13^. We hypothesize that this is also the case for Co^2+^, since *coaT* also encodes an efflux pump. P_*coaT*_ specifically, as well as metal-inducible promoters in general, are unsuitable as inducible promoters in synthetic biology. They lack orthogonality, require laborious alteration of standard culture media, show inducer toxicity at higher concentrations and are outperformed by all three inducible systems shown in this work, both in terms of dynamic range and maximum strength.

### Different inducible promoters cover a wide range of expression levels

Finally, we measured the performance of each promoter alongside each other, either uninduced or induced. In order to categorize each promoter within a broader range, we included the native promoter constructs P_cpc560_, P_*rnpB*_ and P_*rbcL*_, as well as the minimal constitutive promoters P_J23100_ and P_J23119_.

All strains were cultured in accordance with the dose-response assays shown previously. Transconjugants were validated, cultured, and mVenus fluorescence, as well as the optical density at 750 nm, was measured in a microplate reader after 24h. For all four inducible promoters, cultures both uninduced and induced with 10 mM L-rhamnose, 1 μM aTc, 1 mM vanillate or 10 μM CoCl_2_, were grown and measured.

Consistent with the previous results, the fluorescence of P_rha_ is the strongest of the inducible promoters, closely followed by P_L03_ and P_vanCC_ (Fig. 6 A).

**Fig. 6.**
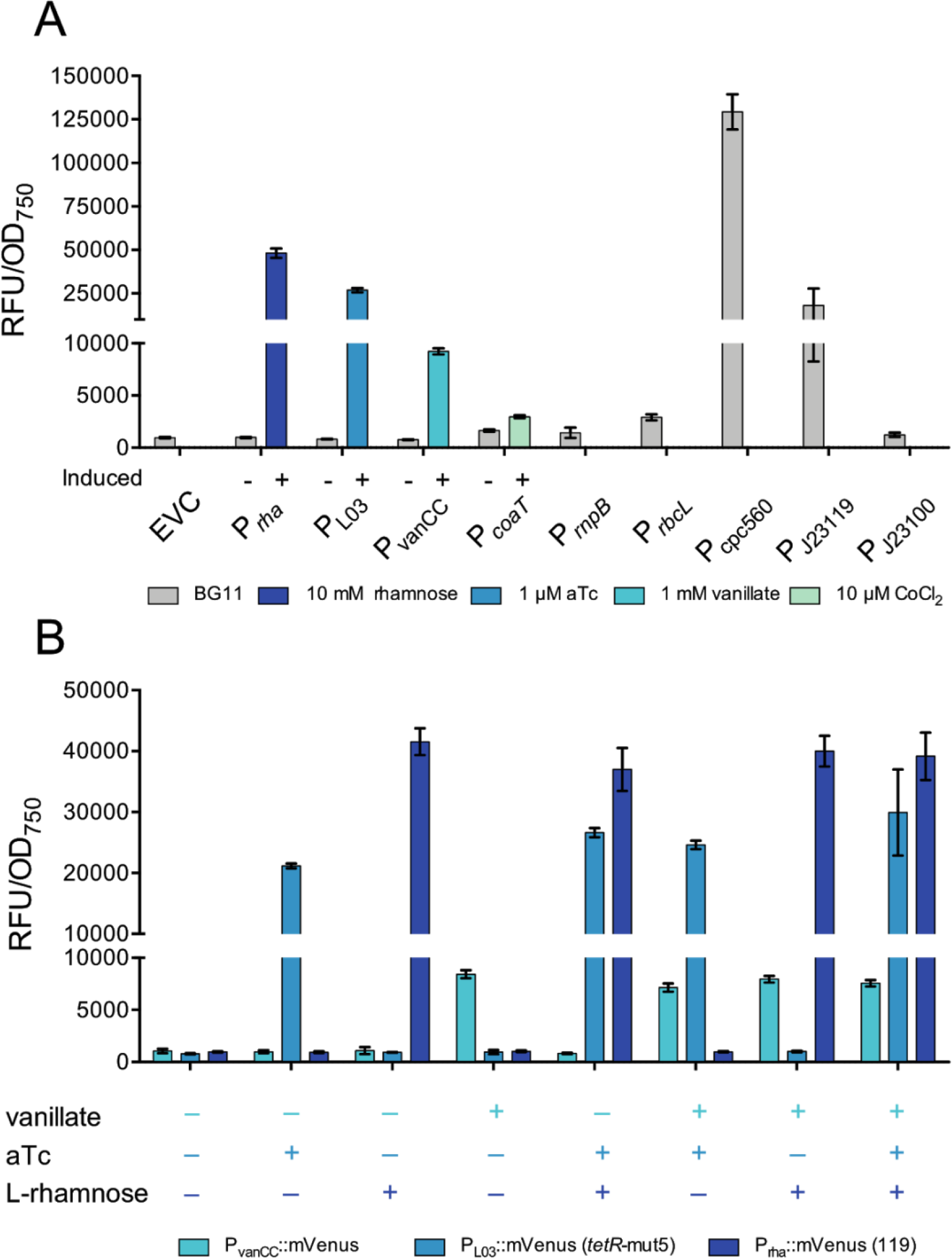
Comparison of established constitutive promoter and inducible promoters. **A:** Comparison of inducible with constitutive promoters. EVC: Empty vector control. P_rha_: P_rha_∷mVenus(119). P_L03_: P_L03_∷mVenus (*tetR*-mut5). **B:** Evaluation of inducer specificity. Three biological replicates each were cultured in BG11 + inducer (10 mM rhamnose, 1 mM vanillate or 1 μM aTc or combinations thereof, marked by a + when present or a − when absent) and fluorescence and optical density was measured in a microplate reader after 24 h.

While these three show promise both in terms of dynamic range and strength, P_*coaT*_ is by far the weakest of the four. The uninduced control, which was cultured in regular BG11 instead of BG11 lacking CoCl_2_, shows leaky induction, leading of a fold change of only 2× for P_*coaT*_.

The strongest inducible promoter, P_rha_, is still weaker than P_cpc560_, the “super-strong” promoter published by Zhou *et al*^31^. This promoter enabled expression of heterologous proteins leading up to 15 % of total soluble protein. However, the data shown in Fig. 6A was measured 24h post-induction, and P_rha_ shows a steady and strong increase in fluorescence over 7 days (Fig. 3D). It could be assumed that P_rha_ is able to reach levels similar to P_cpc560_ after a sufficient induction time.

In order to investigate inducer specificity, each single promoter construct was also induced with all possible combinations of inducers. Cultures were induced with 10 mM L-rhamnose, 1 μM aTc or 1 mM vanillate, or a combination thereof. If left uninduced, the corresponding volume of solvent (H_2_O or ethanol) was added.

All promoters show specific induction only in the presence of the respective inducer molecule (Fig. 6B). The level of fluorescence appears to be the same regardless of the presence or absence of the other inducers for each promoter. In terms of inducer specificity, the promoter constructs are therefore compatible with one another. It remains to be investigated whether they are truly orthogonal to each other in terms of transcription factor binding specificity, i.e., whether the transcriptional regulators are able to bind to unspecific operator sequences and activate or repress gene expression.

## Conclusions & Outlook

In this work, we constructed and evaluated a library of different inducible promoters in a way that enables a useful comparison for later selection of a suitable promoter in *Synechocystis sp*. PCC 6803. Using the pSHDY plasmid allowed efficient exchange of parts to build this library, as well as comparable conditions. We observed a delicate balance between transcription factor toxicity and sufficient expression to obtain a dose-dependent response to the inducer. This observation should be kept in mind for future works, as it might significantly improve the performance of other promoters. Next to the established aTc- and rhamnose-inducible promoters P_L03_ and P_rha_, we report the vanillate inducible promoter P_vanCC_ as a new tool for *Synechocystis*. All three promoters show a linear induction over a range of inducer concentrations, as well as little to no leakiness in the absence of the inducer. Interestingly, they show different strengths of expression, as well as different temporal expression patterns, with the potential for a wide range of biological applications. Thus, our promoter library allows moving away from metal-inducible promoters and towards well-characterized, defined and orthogonal parts, a key requirement of synthetic biology.

The next step in applying the three inducible promoters for future works would be evaluating their performance in a strain genomically encoding the transcriptional regulators. Ultimately, encoding the regulators on the genome using a markerless genomic manipulation strategy would facilitate working with cyanobacteria, since it would free available space on the plasmid, as well as antibiotic resistances. This strategy has proven successful in the past in *E. coli*, resulting in many expression strains for different applications.

Finally, all three promoters should be combined with different reporter genes each and encoded in one strain to evaluate whether they are truly orthogonal and whether they can be used in combination to control multiple genes or operons, enabling the possibility for larger synthetic networks or metabolic engineering by optimization of metabolic pathways.

## Material & Methods

### Plasmid and strain construction

A detailed list of all relevant genetic modules and information regarding their origin, as well as plasmids constructed from them, is provided in the Supplemental Information (Table S1).

All parts were amplified and fused using overlap extension PCR (dx.doi.org/10.17504/protocols.io.psndnde) and integrated into the pSHDY backbone via Gibson assembly (dx.doi.org/10.17504/protocols.io.n9xdh7n).

Plasmids were transferred to *Synechocystis* sp. PCC 6803 wild type using triparental mating (dx.doi.org/10.17504/protocols.io.psndnde). Clones were verified via colony PCR (dx.doi.org/10.17504/protocols.io.mk5c4y6).

### Culture conditions

All strains were maintained on BG11 plates containing 40 μg/mL spectinomycin.

Prior to each assay, BG11 + 20 μg/mL spectinomycin were inoculated with the strain of interest, grown for 5 days, diluted to an OD_750_ of 0.2, grown for 3 more days, and diluted again to the desired OD_750_ (specified in each assay) prior to starting the experiment.

Liquid cultures were grown in constant white light (80 μmol·m^−2^·s^−1^, 16% intensity setting in the Infors HT multitron) at 30 °C and 75% humidity with constant agitation at 150 rpm without added CO_2_.

Detailed protocols for each assay can be found on protocols.io:

Dose response assay → dx.doi.org/10.17504/protocols.io.55wg87e
Toxicity assay → dx.doi.org/10.17504/protocols.io.6tghejw
Fluorescence time course assay → dx.doi.org/10.17504/protocols.io.6tkhekw

### Measurements & settings

To determine cell density, absorbance of cells was measured in a Specord 200 Plus spectrophotometer (Analytik Jena) at 750 nm.

Fluorescence measurements were performed using a BMG Clariostar. Absorbance at 750 nm, as well as fluorescence at λ^ex^/λ^em^ 511/552, was measured every time. Prior to each measurement, the plate was shaken at 500 rpm for 30 seconds.

The exact protocol for the BMG can be found in Supplementary File 1.

### Data analysis & -treatment

For dose response assays, fluorescence values were divided by OD_750_.

For fluorescence time course assays, fluorescence values were divided by OD_750_. Then, the mean of the values measured for the uninduced control culture was subtracted from each individual value measured for the induced culture.

For the fluorescence time courses, all raw fluorescence values were normalized to OD_750_, then, the mean fluorescence of the uninduced control was subtracted from each value of the induced culture.

The plasmid pAJM.714, as well as the strain sAJM.1504 were a gift from Christopher Voigt (Addgene plasmid # 108515; Bacterial strain # 108251).

## Abbreviations

aTc: anhydrotetracycline
*Escherichia coli*: *E. coli*
LAHG: light-activated heterotrophic growth
RBS: ribosome binding site
*Synechocystis*: Synechocystis sp. PCC 6803
WT: wild type
OD_750_: Optical density at 750 nm
TSS: Transcription start site

## Author Contributions

AB and IMA designed and conceived the study. AB and PS performed the experiments and analyzed the data. AB wrote the manuscript with input from PS and IMA.

## References

(1) Oliver, N. J., Rabinovitch-Deere, C. A., Carroll, A. L., Nozzi, N. E., Case, A. E., and Atsumi, S. (2016) Cyanobacterial metabolic engineering for biofuel and chemical production. Current Opinion in Chemical Biology 35, 43–50.

(2) Ducat, D. C., Way, J. C., and Silver, P. A. (2011) Engineering cyanobacteria to generate high-value products. Trends in Biotechnology 29, 95–103.

(3) Zarzycki, J., Axen, S. D., Kinney, J. N., and Kerfeld, C. A. (2013) Cyanobacterial-based approaches to improving photosynthesis in plants. J. Exp. Bot. 64, 787–798.

(4) Ramey, C. J., Barón-Sola, Á., Aucoin, H. R., and Boyle, N. R. (2015) Genome Engineering in Cyanobacteria: Where We Are and Where We Need To Go. ACS Synth. Biol. 4, 1186–1196.

(5) Huang, H.-H., Camsund, D., Lindblad, P., and Heidorn, T. (2010) Design and characterization of molecular tools for a Synthetic Biology approach towards developing cyanobacterial biotechnology. Nucleic Acids Research 38, 2577–2593.

(6) Wang, B., Eckert, C., Maness, P.-C., and Yu, J. (2018) A Genetic Toolbox for Modulating the Expression of Heterologous Genes in the Cyanobacterium Synechocystis sp. PCC 6803. ACS Synth. Biol. 7, 276–286.

(7) Kim, W. J., Lee, S.-M., Um, Y., Sim, S. J., and Woo, H. M. (2017) Development of SyneBrick Vectors As a Synthetic Biology Platform for Gene Expression in Synechococcus elongatus PCC 7942. Front. Plant Sci 8, 1–9.

(8) Santos-Merino, M., Singh, A. K., and Ducat, D. C. (2019) New Applications of Synthetic Biology Tools for Cyanobacterial Metabolic Engineering. Front. Bioeng. Biotechnol. 7, 273.

(9) Higo, A., and Ehira, S. (2019) Spatiotemporal Gene Repression System in the Heterocyst-Forming Multicellular Cyanobacterium Anabaena sp. PCC 7120. ACS Synth. Biol. 8, 641–646.

(10) Vasudevan, R., Gale, G. A. R., Schiavon, A. A., Puzorjov, A., Malin, J., Gillespie, M. D., Vavitsas, K., Zulkower, V., Wang, B., Howe, C. J., Lea-Smith, D. J., and McCormick, A. J. (2019) CyanoGate: A Modular Cloning Suite for Engineering Cyanobacteria Based on the Plant MoClo Syntax. Plant Physiology 180, 39–55.

(11) Markley, A. L., Begemann, M. B., Clarke, R. E., Gordon, G. C., and Pfleger, B. F. (2014) Synthetic Biology Toolbox for Controlling Gene Expression in the Cyanobacterium Synechococcus sp. strain PCC 7002. ACS Synth. Biol. 4, 595–603.

(12) Imamura, S., and Asayama, M. (2009) Sigma Factors for Cyanobacterial Transcription. Gene Regulation and Systems Biology 1–23.

(13) Englund, E., Liang, F., and Lindberg, P. (2016) Evaluation of promoters and ribosome binding sites for biotechnological applications in the unicellular cyanobacterium Synechocystis sp. PCC 6803. Sci. Rep. 6, 36640.

(14) Geerts, D., Bovy, A., De Vrieze, G., Borrias, M., and Weisbeek, P. (1995) Inducible expression of heterologous genes targeted to a chromosomal platform in the cyanobacterium Synechococcus sp. PCC 7942. Microbiology 141, 831–841.

(15) Ferreira, E. A., Pacheco, C. C., Pinto, F., Pereira, J., 2018. Expanding the toolbox for Synechocystis sp. PCC 6803: validation of replicative vectors and characterization of a novel set of promoters. Synthetic Biology.

(16) Camsund, D., Heidorn, T., and Lindblad, P. (2014) Design and analysis of LacI-repressed promoters and DNA-looping in a cyanobacterium. Journal of Biological Engineering 8, 1–23.

(17) Albers, S. C., Gallegos, V. A., and Peebles, C. A. M. (2015) Engineering of genetic control tools in Synechocystis sp. PCC 6803 using rational design techniques. Journal of Biotechnology 216, 36–46.

(18) Huang, H.-H., and Lindblad, P. (2013) Wide-dynamic-range promoters engineered for cyanobacteria. Journal of Biological Engineering 7, 1–11.

(19) Yao, L., Cengic, I., Anfelt, J., and Hudson, E. P. (2016) Multiple Gene Repression in Cyanobacteria Using CRISPRi. ACS Synth. Biol. 5, 207–212.

(20) Kelly, C. L., Taylor, G. M., Hitchcock, A., Torres-Méndez, A., and Heap, J. T. (2018) A Rhamnose-Inducible System for Precise and Temporal Control of Gene Expression in Cyanobacteria. ACS Synth. Biol. 7, 1056–1066.

(21) Thiel, K., Mulaku, E., Dandapani, H., Nagy, C., Aro, E.-M., and Kallio, P. (2018) Translation efficiency of heterologous proteins is significantly affected by the genetic context of RBS sequences in engineered cyanobacterium Synechocystis sp. PCC 6803. Microbial Cell Factories 17, 1–12.

(22) Cho, S. H., Haning, K., Shen, W., Blome, C., Li, R., Yang, S., and Contreras, L. M. (2017) Identification and Characterization of 5’ Untranslated Regions (5’UTRs) in Zymomonas mobilis as Regulatory Biological Parts. Front. Microbiol. 8, 233.

(23) Los, D. A., Zorina, A., Sinetova, M., Kryazhov, S., Mironov, K., and Zinchenko, V. V. (2010) Stress Sensors and Signal Transducers in Cyanobacteria. Sensors 10, 2386–2415.

(24) Ruegg, T. L., Pereira, J. H., Chen, J. C., Nature, A. D., 2018. Jungle Express is a versatile repressor system for tight transcriptional control. Nature Communications.

(25) Meyer, A. J., Segall-Shapiro, T. H., Glassey, E., Zhang, J., and Voigt, C. A. (2019) Escherichia coli “Marionette” strains with 12 highly optimized small-molecule sensors. Nature Chemical Biology 15, 196–204.

(26) Zinchenko, V. V., Piven, I. V., Melnik, V. A., and Shestakov, S. V. (1999) Vectors for the Complementation Analysis of Cyanobacterial Mutants. Russian Journal of Genetics 35, 228–232.

(27) Taton, A., Unglaub, F., Wright, N. E., Zeng, W. Y., Paz-Yepes, J., Brahamsha, B., Palenik, B., Peterson, T. C., Haerizadeh, F., Golden, S. S., and Golden, J. W. (2014) Broad-host-range vector system for synthetic biology and biotechnology in cyanobacteria. Nucleic Acids Research 42, e136–e136.

(28) Kremers, G.-J., Goedhart, J., van Munster, E. B., and Gadella, T. W. J. (2006) Cyan and yellow super fluorescent proteins with improved brightness, protein folding, and FRET Förster radius. Biochemistry 45, 6570–6580.

(29) Andersen, J. B., Sternberg, C., Poulsen, L. K., Bjorn, S. P., Givskov, M., and Molin, S. (1998) New unstable variants of green fluorescent protein for studies of transient gene expression in bacteria. Applied and Environmental Microbiology 64, 2240–2246.

(30) Heidorn, T., Camsund, D., Huang, H.-H., Lindberg, P., Oliveira, P., Stensjö, K., and Lindblad, P. (2011) Synthetic biology in cyanobacteria engineering and analyzing novel functions. Methods in Enzymology 497, 539–579.

(31) Zhou, J., Zhang, H., Meng, H., Zhu, Y., Bao, G., Zhang, Y., Li, Y., and Ma, Y. (2014) Discovery of a super-strong promoter enables efficient production of heterologous proteins in cyanobacteria. Sci. Rep. 4, 235–6.

(32) Rawlings, D. E., and Tietze, E. (2001) Comparative Biology of IncQ and IncQ-Like Plasmids. Microbiology and Molecular Biology Reviews 65, 481–496.

(33) Kaczmarczyk, A., Vorholt, J. A., and Francez-Charlot, A. (2014) Synthetic vanillate-regulated promoter for graded gene expression in *Sphingomonas*. Sci. Rep. 4, 6453.

(34) Taton, A., Ma, A. T., Ota, M., Golden, S. S., and Golden, J. W. (2017) NOT Gate Genetic Circuits to Control Gene Expression in Cyanobacteria. ACS Synth. Biol. 6, 2175–2182.

(35) Kamimura, N., Takahashi, K., Mori, K., Araki, T., Fujita, M., Higuchi, Y., and Masai, E. (2017) Bacterial catabolism of lignin-derived aromatics: New findings in a recent decade: Update on bacterial lignin catabolism. Environmental Microbiology Reports 9, 679–705.

(36) Oliva, B., Gordon, G., McNicholas, P., Ellestad, G., and Chopra, I. (1992) Evidence that tetracycline analogs whose primary target is not the bacterial ribosome cause lysis of Escherichia coli. Antimicrobial Agents and Chemotherapy 36, 913–919.

(37) Mathieu, A., Fleurier, S., Frénoy, A., Dairou, J., Bredeche, M.-F., Sanchez-Vizuete, P., Song, X., and Matic, I. (2016) Discovery and Function of a General Core Hormetic Stress Response in E. coli Induced by Sublethal Concentrations of Antibiotics. CellReports 17, 46–57.

(38) Loeschcke, A., Dienst, D., Wewer, V., Hage-Hülsmann, J., Dietsch, M., Kranz-Finger, S., Hüren, V., Metzger, S., Urlacher, V. B., Gigolashvili, T., Kopriva, S., Axmann, I. M., Drepper, T., and Jaeger, K.-E. (2017) The photosynthetic bacteria Rhodobacter capsulatus and Synechocystis sp. PCC 6803 as new hosts for cyclic plant triterpene biosynthesis. PLoS ONE 12, e0189816.

(39) Guerrero, F., Carbonell, V., Cossu, M., Correddu, D., and Jones, P. R. (2012) Ethylene Synthesis and Regulated Expression of Recombinant Protein in Synechocystis sp. PCC 6803. PLoS ONE (Neilan, B., Ed.) 7, e50470–11.

(40) González, A., Bes, M. T., Peleato, M. L., and Fillat, M. F. (2016) Expanding the Role of FurA as Essential Global Regulator in Cyanobacteria. PLoS ONE (Hess, W. R., Ed.) 11, e0151384.

(41) Cavet, J. S., Borrelly, G. P. M., and Robinson, N. J. (2003) Zn, Cu and Co in cyanobacteria: selective control of metal availability. FEMS Microbiol Rev 27, 165–181.

(42) Berla, B. M., Saha, R., Immethun, C. M., Maranas, C. D., Moon, T. S., and Pakrasi, H. (2013) Synthetic biology of cyanobacteria: unique challenges and opportunities. Front. Microbiol. 4.

(43) Holm Hansen, O., Gerloff, G. C., and Skoog, F. (1954) Cobalt as an Essential Element for Blue-Green Algae. Physiologia Plantarum 7, 665–675.

