## Supplementary File 1 for "Comparative analysis of inducible promoters in cyanobacteria"

;Script generated by script wizard (CLARIOstar V5.60 R2)
;Thomas Seidel, BMG LABTECH, January 2013 ... January 2019

;Protocol Names:
st1:="abs_750_Anna"
st2:="mVenus_511_552_Anna"

;Plate ID:
ID1:="<date>,<time>, "

;Number of readings (number of kinetic cycles):
NumberOfReadings:=1

;Define the time in seconds after which the plate should be measured again:
CycleTime:=60

;==============================================================================================

st9:=", <date> <time>"  ;part of plate ID 2

ID2:="Wizard script 'Anna_Cyano'<st9>"

ID3:="unmerged"  ;marker for Merge tool

R_PlateUser 0 0                                                                          ;move to defined start position

for Reading:=1 to NumberOfReadings do begin                          ;kinetic loop
    CycleStartTime:=<now>

    R_Run "<st1>"                                                                            ;execute first test protocol

    R_Run "<st2>"                                                                            ;execute second test protocol

    ;merge vertical data from first two test runs:
    Call "MergeReadings.exe <DataPath> <User> VU"

    ;merge horizontal (kinetic):
    if Reading>1 then begin
        Call "MergeReadings.exe <DataPath> <User> H ID2"
    end;

    ;Wait until the defined cycle time is over:
    if Reading<NumberOfReadings then begin                                 ;Waiting is not necessary after the last measurement!
        R_PlateUser 0 0                                                                      ;move to defined start position

        MeasurementEndTime:=<now>
        CycleDurationSoFar:=MeasurementEndTime-CycleStartTime
        CycleDurationSoFar:=86400*CycleDurationSoFar                   ;convert fractions of a day into seconds
        RemainingCycleTime:=CycleTime-CycleDurationSoFar           ;remaining time to reach the defined cycle time

        if RemainingCycleTime>0 then begin
            wait for RemainingCycleTime s                                            ;wait remaining time
         end;
     end;
end;                                                                                             ;end of kinetic loop

;end of script
