## Supplementary File 2 for "Comparative analysis of inducible promoters in cyanobacteria"

Supporting Information

Supplementary File 1: BMG ClarioStar protocol for measurement of mVenus fluorescence in *Synechocystis* sp. PCC 6803

Supplementary File 2: Supporting Tables and Figures for this work.

Table S1: Detailed descriptions and sequences of all relevant genetic modules used in this work

Fig. S1: 5'RACE results from TSS-mapping of $P_{coaT}$.

To map the transcription start site (TSS) of $P_{coaT}$, we performed 5'RACE, as described in detail on protocols.io (dx.doi.org/10.17504/protocols.io.jk7ckzn). Five putative TSS were discovered, three of which occurred at a higher frequency (Fig. S1).

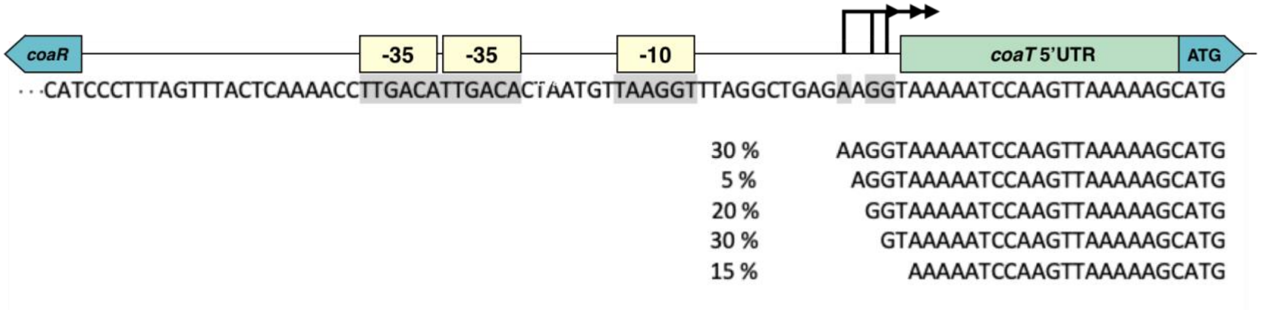


**Fig. S1: 5'RACE results from TSS-mapping of $P_{coaT}$.**
Top: Schematic overview of genetic construct used. -10, -35 and putative +1 are highlighted in grey.
Bottom: Putative transcription start sites based on sequencing results from 5'RACE. The percentages indicate the frequency of each individual start site as determined by sequencing. A total of 20 clones were sequenced.

**Table S1: Detailed descriptions and sequences of all relevant genetic modules used in this work.**

| Name | Part type | Sequence | Origin | Notes |
|---|---|---|---|---|
| $P_{vanCC}$::riboJ | Promoter + insulator | gattggatccaattgacagctagctcagtcctaggtaccattggatccaatagctgtcaccggatgtgctttccggtctgatgagtccgtgaggacgaaacagcctctacaaataattttgtttaa | [25] | Amplified from addgene plasmid pAJM.714 |
| RBS* | RBS | tactagagtagtggaggttactag | [30] | |
| mVenus_ASV | CDS | atggctagcaaaggagaggaactgtttaccggtgtggtacccattttagtagaattggatggtgatgtgaacggccacaagttcagcgtttccggggaaggcgaaggggatgcaacctatggaaagctaactttgaaactcatttgcactaccggtaaactccccgtcccctggcctacgttagtcacgaccctcggttatggtctgcaatgttttgcccgttatcccgatcatatgaaacaacacgatttctttaaatctgccatgccagagggatatgttcaagaacggacaattttctttaaggatgatggtaattataaaacccgtgcagaagtcaaatttgaaggtgatacactcgtgaatcgcatcgaactaaaaggtattgactttaaggaggatggaaacattctcggccataagttagaatacaattacaacagccataatgtgtacattactgcggataaacaaaaaacggtatcaaagccaactttaaaattcgacataacatcgaagatggtggcgttcagttagcggatcactaccaacaaaatactcccattggagacggaccagtcctgctgcctgataaccattatctgtcttatcaatccaaactgtccaaggatccgaatgaaaagcgggatcacatggttctgttggagtttgtgaccgcggcaggtattactctcggaatggacgaactatacaaaaggcctgccgccaacgacgaaaactacgctgcatcagtttaa | This work | Codon-optimized for *Synechocystis*, with C-terminal ASV degradation tag (highlighted in grey). |
| $P_{rha}$ | Promoter | gccacaattcagcaaattgtgaacatcatcacgttcatctttccctggttgccaatggcccattttcctgtcagtaacgagaaggtcgcgaattcaggcgctttttagactggtcgtaatgaa | [20] | |
| $P_{L03}$ | Promoter | gtacgcaattggcatgcgcattccctatcagtgatagagattgacatccctatcagtgatagatataatggccact | [18] | |
| $P_{coaT}$ | Promoter +5'UTR | ccctttagtttactcaaaaccttgacattgacactaatgttaaggtttaggctgagaaggtaaaaatccaagttaaaaagc | *Synechocystis* | 81 bp upstream of the *coaT* gene |
| RBSvan3::*vanR* | RBS + CDS | gcttaaactaacgaacgtaaataaggaggatagacatggacatgcctcgtattaaaccgggtcagcgtgttatgatggcactgcgtaaaatgattgcaagcggtgaaatcaaaagtggtgaacgtattgcagaaattccgaccgcagcagcactgggtgttagccgtatgccggttcgtatcgcactgcgttcactggaacaagaaggtctggttgttcgtctgggtgcacgtggttatgcagcccgtggtgttagcagcgatcagattcgtgatgcaattgaagttcgtggtgttctggaaggttttgcagcacgtcgtctggcagaacgtggtatgaccgcagaaacccatgcacgttttgttgtactgattgcagaaggtgaagcactgtttgcagccggtcgcctgaatggtgaagatctggatcgttatgccgcatataatcaggcatttcatgataccctggttagcgcagcaggtaatggtgcagttgaaagcgcactggcacgtaatggttttgaaccgtttgcagcagccggtgcactggccctggatctgatggacctgtctgccgaatatgaacatctgctggcagcacatcgtcagcatcaggcagttctggatgcagttagctgtggtgatgccgaaggtgcagaacgtattatgcgtgatcatgcactggcagcaattcgtaatgcaaaagtttttgaagcagcagcaagcgcaggcgcaccgctgggtgcagcatggtcaattcgtgcagattga | [25] | Amplified from addgene strain sAJM.1504 |
| $P_{J23119}$ | Promoter | ttgacagctagctcagtcctaggtataatgctagc | iGEM registry | Bba_J23119 |
| $P_{J23111}$ | Promoter | ttgacggctagctcagtcctaggtatagtgctagc | iGEM registry | Bba_J23119 |
| $P_{J23100}$ | Promoter | ttgacggctagctcagtcctaggtacagtgctagc | iGEM registry | Bba_J23119 |
| Bba_0034 | RBS | aaagaggagaaatactag | iGEM registry | Bba_B0034 |
| *rhaS* | CDS | atgaccgtattacatagtgtggatttttttccgtctggtaacgcgtccgtggcgatagaaccccggctcccgcaggcggattttcctgaacatcatcatgattttcatgaaattgtgattgtcgaacatggcacgggtattcatgtgtttaatgggcagccctataccatcaccggtggcacggtctgtttcgtacgcgatcatgatcggcatctgtatgaacataccgataatctgtgtctgaccaatgtgctgtatcgctcgccggatcgatttcagtttctcgccgggctgaatcagttgctgccacaagagctggatgggcagtatccgtctcactggcgcgttaaccacagcgtattgcagcaggtgcgacagctggttgcacagatggaacagcaggaaggggaaaatgatttaccctcgaccgccagtcgcgagatcttgtttatgcaattactgctcttgctgcgtaaaagcagtttgcaggagaacctggaaaacagcgcatcacgtctcaacttgcttctggcctggctggaggaccattttgccgatgaggtgaattgggatgccgtggcggatcaattttctctttcactgcgtacgctacatcggcagcttaagcagcaaacgggactgacgcctcagcgatacctgaaccgcctgcgactgatgaaagcccgacatctgctacgccacagcgaggccagcgttactgacatcgcctatcgctgtggattcagcgacagtaaccacttttcgacgctttttcgccgagagtttaactggtcaccgcgtgatattcgccagggacgggatggctttctgcaataa | *E. coli* | Amplified from *E. coli* genome |
| $P_{J23119}$-mut5 | Promoter + RBS | ttgacagctagctcagtcctaggtataatgctagcaagagagaaatactag | this work | |
| $P_{J23119}$-mut5 | Promoter + RBS | ttaacagctagctcagtcctaggtataatgctagcaaagaggagaaatactag | this work | |
| *tetR* | CDS | atgtccagattagataaaagtaaagtgattaacagcgcattagagctgcttaatgaggtcggaatcgaaggtttaacaacccgtaaactcgcccagaagctaggtgtagagcagcctacattgtattggcatgtaaaaaataagcgggctttgctcgacgccttagccattgagatgttagataggcaccatactcacttttgccctttagaaggggaaagctggcaagattttttacgtaataacgctaaaagttttagatgtgctttactaagtcatcgcgatggagcaaaagtacatttaggtacacggcctacagaaaaacagtatgaaactctcgaaaatcaattagcctttttatgccaacaaggtttttcactagagaatgcattatatgcactcagcgctgtggggcattttactttaggttgcgtattggaagatcaagagcatcaagtcgctaaagaagaaagggaaacacctactactgatagtatgccgccattattacgacaagctatcgaattatttgatcaccaaggtgcagagccagccttcttattcggccttgaattgatcatatgcggattagaaaaacaacttaaatgtgaaagtgggtcctaa | iGEM registry | Bba_P0440. *tetR* was amplified omitting the LVA degradation tag. |
| $P_{rnpB}$ | Promoter | ttcaatgcggtccaatacctcccctgcccaactgggtaagctcgcggctccactgagtaatacagacaaggctaaacaggcaaattttttcattggtcaactcctagcaccaatttcccaagactacggagggggcaatgaagtttcaattaattggggtcacaaaccacagcggcctatggctctaatcaatggcacactagaaaaa | *Synechocystis* | Amplified from *Synechocystis* genome |
| $P_{rbcL}$ | Promoter + 5'UTR | cagtcaatggagagcattgccataagtaaaggcatcccctgcgtgataagattaccttcagaaaacagatagttgctgggttatcgcagatttttctcgcaaccaaataactgtaaataataactgtctctggggcgacggtaggctttatattgccaaatttcgcccgtgggagaaagctaggctatt | *Synechocystis* | Amplified from *Synechocystis* genome |
| $P_{cpc560}$ | Promoter + 5'UTR | cacctgtagagaagagtccctgaatatcaaaatggtgggataaaaagctcaaaaaggaaagtaggctgtggttccctaggcaacagtcttccctaccccactggaaactaaaaaaacgagaaaagttcgcaccgaacatcaattgcataattttagccctaaaacataagctgaacgaaactggttgtcttcccttcccaatccaggacaatctgagaatcccctgcaacattacttaacaaaaaagcaggaataaaattaacaagatgtaacagacataagtcccatcaccgttgtataaagttaactgtgggattgcaaaagcattcaagcctaggcgctgagctgtttgagcatcccggtggcccttgtcgctgcctccgtgtttctccctggatttatttaggtaatatctctcataaatccccgggtagttaacgaaagttaatggagatcagtaacaataactctagggtcattactttggactccctcagtttatccgggggaattgtgtttaagaaaatcccaactcataaagtcaagtaggagattaattca | *Synechocystis* [31] | Amplified from *Synechocystis* genome |
